# A truncated form of HpARI stabilises IL-33, amplifying responses to the cytokine

**DOI:** 10.1101/2020.04.13.035477

**Authors:** Caroline Chauché, Francesco Vacca, Shin Li Chia, Josh Richards, William F Gregory, Adefunke Ogunkanbi, Martin Wear, Henry J McSorley

## Abstract

The murine intestinal nematode *Heligmosomoides polygyrus* releases the *H. polygyrus* Alarmin Release Inhibitor (HpARI) - a protein which binds to IL-33 and to DNA, effectively tethering the cytokine in the nucleus of necrotic cells. Previous work showed that a non-natural truncation consisting of the first 2 domains of HpARI (HpARI_CCP1/2) retains binding to both DNA and IL-33, and inhibited IL-33 release *in vivo*. Here, we show that the affinity of HpARI_CCP1/2 for IL-33 is significantly lower than that of the full-length protein, and that HpARI_CCP1/2 lacks the ability to prevent interaction of IL-33 with its receptor. When HpARI_CCP1/2 was applied *in vivo* it potently amplified IL-33-dependent immune responses to *Alternaria alternata* allergen, *Nippostrongylus brasiliensis* infection and recombinant IL-33 injection. Mechanistically, we found that HpARI_CCP1/2 is able to bind to and stabilise IL-33, preventing its degradation and maintaining the cytokine in its active form. This study highlights the importance of IL-33 inactivation, the potential for IL-33 stabilisation *in vivo*, and describes a new tool for IL-33 research.

## 2 Introduction

*Heligmosomoides polygyrus* is a parasitic nematode that infects the intestines of mice. It has a faecal/oral lifecycle, with infective L3 larvae being ingested, and then rapidly penetrating the epithelium of the proximal duodenum. There, the larvae develop to L4 stage and emerge as adults into the intestinal lumen at around day 10 of infection (Johnston et al., 2015; Reynolds, Filbey, & Maizels, 2012). The transit of the parasite through the intestinal wall is likely to cause epithelial damage and cell death, with resulting release of alarmins such as IL-33 (from stromal cells or mast cells (Shimokawa et al., 2017)), in turn inducing an anti-parasite type 2 immune response (Harris & Loke, 2017). In order to negate this response, and allow persistence of the parasite in the host, *H. polygyrus* secretes multiple immunomodulatory factors, including Hp-TGM, a protein mimic of host TGF-β (Johnston et al., 2017), and microRNA-containing extracellular vesicles (Buck et al., 2014) which modulate transcription of multiple host genes, including suppression of Suppression of Tumorigenicity 2 (ST2), the IL-33 receptor. We previously showed that the parasite also secretes the *H. polygyrus* Alarmin Release Inhibitor (HpARI), which blocks IL-33 responses (Osbourn et al., 2017).

IL-33 is an alarmin cytokine constitutively produced by epithelial cells. It is stored preformed in the nucleus and released on necrotic cell death, due to mechanical, protease-mediated or chemical damage to the epithelium (Johansson & McSorley, 2019). On necrotic cell death, proteases from the cell cytoplasm, or those secreted by recruited mast cells, neutrophils or those in allergens can then cleave the cytokine between the N-terminus chromatin-binding domain and the C-terminus receptor binding domain, potently increasing the activity of the cytokine (Cayrol et al., 2018; Lefrancais et al., 2012; Scott et al., 2018). The IL-33 receptor-binding domain contains four free cysteine residues, which upon release from the reducing nuclear environment into the oxidising extracellular environment rapidly form disulphide bonds, changing the cytokine’s conformation, rendering it unable to bind to its receptor and effectively inactivating it (Cohen et al., 2015). Proteases can also further degrade IL-33 to smaller, inactive forms (Scott et al., 2018). Thus, the active form of IL-33 has only a very short half-life, and by 1 hour after release the vast majority of IL-33 is inactive or degraded.

HpARI binds to the active reduced form of IL-33 and to genomic DNA. This dual binding tethers IL-33 within the nucleus of necrotic cells, preventing its release, and inhibiting interaction of IL-33 with ST2. The HpARI protein consists of 3 Complement Control Protein domains (CCP1-3), and our previous data showed that HpARI binds IL-33 through the CCP2 domain, while DNA-binding was mediated by the CCP1 domain (Osbourn et al., 2017). Here, we further characterise the functions of the CCP domains of HpARI, finding that CCP3 stabilises the interaction between HpARI and IL-33, increasing its affinity and being required for blockade of IL-33-ST2 interactions. Furthermore, we show that HpARI_CCP1/2 (the HpARI truncation lacking CCP3) is able to stabilise IL-33, increasing its half-life and amplifying its effects.

## 3 Materials and Methods

### Protein expression and purification

Constructs encoding HpARI, HpARI_CCP1/2 and HpARI_CCP2/3 (all with C-terminus myc and 6-His tags) were cloned into the pSecTAG2A expression vector as previously described (Osbourn et al., 2017). Purified plasmids were transfected into Expi293F™ cells, and supernatants collected 5 days later. Expi293F™ cells were maintained, and transfections carried out using the Expi293 Expression System according to manufacturer’s instructions (ThermoFisher Scientific). Expressed protein in supernatants were purified over a HisTrap excel column (GE Healthcare) and eluted in 500 mM imidazole. Eluted protein was then dialysed to PBS, and repurified on a HiTRAP chelating HP column (GE Healthcare) charged with 0.1 M NiSO4. Elution was performed using an imidazole gradient and fractions positive for the protein of interest were pooled, dialysed to PBS and filter-sterilised. Protein concentration was measured at A280 nM (Nanodrop, ThermoFisher Scientific), using calculated extinction coefficient.

### Surface Plasmon Resonance (SPR)

SPR measurements were performed using a BIAcore T200 instrument (GE Healthcare). Ni^2+^- nitrilotriacetic acid (NTA) sensor chips, 1-ethyl-3-(3-diaminopropyl) carbodiimide hydrochloride (EDC), *N*-hydroxysuccinimide (NHS) and ethanolamine (H_2_N(CH_2_)_2_OH) were purchased from GE Healthcare. HpARI, HpARI_CCP1/2 or HpARI_CCP2/3 were immobilized and covalently stabilised on an NTA sensor chip essentially as described (Wear & Walkinshaw, 2006) with the following modifications: following Ni^2+^ priming (30 sec injection of 500 μM NiCl_2_ at 5 μl·min^-1^), dextran surface carboxylate groups were minimally activated by an injection of 0.2 M EDC; 50 mM NHS at 5 μl·min^-1^ for 240 sec. Respective proteins (at concentrations between 10 and 400 nM), in 10 mM NaH_2_PO_4_, pH 7.5; 150 mM NaCl; 50 μM EDTA; 0.05% surfactant P20, were captured *via* the hexa-his tag and simultaneously covalently stabilised to 400 RU, by varying the contact time. Immediately following the capture/stabilisation a single 15 sec injection of 350 mM EDTA and 50 mM Imidazole in 10 mM NaH_2_PO_4_, pH 7.5; 150 mM NaCl; 50 mM EDTA; 0.05% surfactant P20, at 30 μl·min^-1^, was used to remove non-covalently bound protein, followed by a 180 sec injection of 1 M H2N(CH2)2OH, pH 8.5 at 5 μl·min^-1^. Prior to any experiments, the surface was further conditioned with a 600 sec wash with 10 mM NaH_2_PO_4_, pH 7.5; 150 mM NaCl; 50 μM EDTA; 0.05% surfactant P20 at 100 μl·min^-1^.

SPR single-cycle kinetic titration binding experiments were performed at 25°C. Three-fold dilution series of mIL-33 (2.47 nM to 200 nM), were injected over the sensor surface, in 10 mM NaH_2_PO_4_, pH 7.5; 150 mM NaCl; 50 μM EDTA; 0.05% surfactant P20, at 30 ml.min^-1^ for 30 s followed by a final 600 s dissociation phase. The on- (*k*_+_) and off-rate (*k*_−_) constants and the equilibrium dissociation constants were calculated from the double referenced sensorgrams by global fitting of a 1:1 binding model, with mass transport considerations, using analysis software (v2.02) provided with the Biacore T200 instrument.

### Immunoprecipitation

Protein G dynabeads (ThermoFisher Scientific) were coated with 1 μg mouse ST2-Fc (Biolegend), and washed on a DynaMag-2 magnet with PBS 0.02% Tween 20. 100 ng recombinant murine IL-33 (Biolegend) was then mixed with 1 μg HpARI, HpARI_CCP1/2 or HpARI_CCP2/3, and incubated at room temperature for 15 min, prior to adding to ST2-Fc-coated protein G dynabeads. Beads were washed and bound IL-33 eluted with 50 mM glycine pH2.8, then ran on 4-12% SDS-PAGE gels (ThermoFisher Scientific) under reducing conditions, and transferred to nitrocellulose membranes for western blotting, probing with anti-IL-33 goat polyclonal antibody (R&D Systems AF3626), rabbit anti-goat IgG-HRP secondary antibody (ThermoFisher Scientific) and detected using WesternSure Premium reagent (Licor).

### Animals

BALB/cAnNCrl and C57BL/6JCrl mice were purchased from Charles River, UK. Heterozygous IL-13eGFP^+/GFP^ mice (Neill et al., 2010) were bred in-house. All mice were accommodated and procedures performed under UK Home Office licenses with institutional oversight performed by qualified veterinarians.

### Alternaria models

*Alternaria* allergen was used *in vivo* as previously described (McSorley, Blair, Smith, McKenzie, & Maizels, 2014; Osbourn et al., 2017). *Alternaria* allergen (10 μg), OVA (20 μg), HpARI (10 μg) and HpARI_CCP1/2 (10 μg) were intranasally administered to BALB/c mice. Where indicated, the OVA-specific response was recalled by daily intranasal administration of 20 μg OVA protein on days 14, 15 and 16. Tissues were harvested 24 h or 17 days after initial *Alternaria* allergen administration. Lungs were flushed with 4 washes of 0.5 ml ice-cold PBS to collect bronchoalveolar lavage cells, followed by lung dissection for single cell preparation.

### *Nippostrongylus brasiliensis* infection

The life cycle of *N. brasiliensis* was maintained in Sprague-Dawley rats as previously described (Lawrence, Gray, Osborne, & Maizels, 1996), and infective L3 larvae were prepared from 1-3 week rat fecal cultures. BALB/c mice were subcutaneously infected with 400 L3 *N. brasiliensis* larvae, and culled 3 or 6 days later.

### Intraperitoneal IL-33 treatment

Recombinant murine IL-33 (Biolegend) was injected intraperitoneally to C57BL/6 mice (100 ng/mouse). Mice were culled 3 hours later and peritoneal lavage cells collected in 3 washes of 3 ml ice-cold RPMI.

### Flow cytometry

Cells were stained with Fixable Blue Live/Dead stain (ThermoFisher Scientific), then blocked with anti-mouse CD16/32 antibody and surface stained with CD3 (FITC, clone 145-2C11), CD5 (FITC, clone 53-7.3), CD11b (FITC, M1/70), CD19 (FITC, clone 6D5), GR1 (FITC, clone RB6-8C5), CD45 (AF700, clone 30-F11), ICOS (PCP, clone 15F9), CD4 (PE-Dazzle, cloneRM4.5), CD11c (AF647, clone N418), Ly6G (PerCP, clone 1A8), CD25 (BV650, clone PC61) (Biolegend), CD49b (FITC, clone DX5), ST2 (APC, clone RMST2-2) (ThermoFisher Scientific), Siglec-F (PE, clone ES22-10D8) (Miltenyi). The lineage stain consisted of CD3, CD5, CD11b, CD19, CD49b and GR1, all on FITC. Samples were acquired on an LSR Fortessa (BD Biosciences) and analysed using FlowJo 10 (Treestar).

### CMT-64 cell line

CMT-64 cells (ECACC 10032301) were maintained by serial passage in “complete” RPMI (RPMI 1640 medium containing 10% fetal bovine serum, 2 mM L-glutamine, 100 U/ml Penicillin and 100 μg/ml Streptomycin (ThermoFisher Scientific)) at 37°C, 5% CO2. Cells were seeded into 24- or 96-well plates for Triton-X100 or freeze-thaw treatment respectively. Cells were grown to 100% confluency prior to 2 washes with PBS. For Triton-X100 treatment, cells were then washed into RPMI 1640 containing 0.1% BSA with or without 0.1% Triton-X100, and incubated at 37°C as indicated, prior to collection of supernatants and measurement of IL-33 by ELISA and western blot. For freeze-thaw assays, cells were then washed into complete RPMI containing 10 μg/ml of HpARI or HpARI_CCP1/2, frozen on dry ice for at least 1 h, then thawed and incubated at 37°C as indicated, prior to collection of supernatants and application to bone marrow cell cultures.

### Cytokine measurement

ELISAs were carried out to manufacturer’s instructions for IL-5, IL-13 (Ready-SET-go, ThermoFisher Scientific) and IL-33 (Duoset, Biotechne). IL-33 was also measured in CMT-64 supernatants by western blot – supernatants were ran on 4-12% NuPAGE gels (ThermoFisher Scientific) under reducing conditions, before transferring to nitrocellulose membrane and probing with goat anti-mIL-33 (Biotechne), and rabbit anti-goat IgG HRP secondary antibody (Thermo Fisher), and detected using WesternSure Premium reagent (Licor).

### Statistical Analysis

All data was analysed using Prism (Graphpad Software Inc.). One-way ANOVA with Dunnet’s multiple comparisons post-test was used to compare multiple independent groups, while two-way ANOVA and Tukey’s multiple comparison’s post-test was used to compare multiple timepoints or concentrations between independent groups. Where necessary, data was log-transformed to give a normal distribution and to equalise variances. **** = p<0.0001, *** = p<0.001, ** = p<0.01, * = p<0.05, N.S. = Not Significant (p>0.05).

## 4 Results

### HpARI CCP2 binds IL-33, while HpARI CCP3 is required to block IL-33-ST2 interaction

Constructs encoding full-length HpARI, or truncations lacking CCP3 (HpARI_CCP1/2), or lacking CCP1 (HpARI_CCP2/3) were expressed in Expi293F^™^ mammalian cells, and purified on 6-His tags. These constructs were then tested for binding to IL-33 in surface plasmon resonance experiments, showing that the affinity for IL-33 of full-length HpARI and HpARI_CCP2/3 were similar (Kd of 1.1+/-0.44 nM and 1.4 +/- 0.14 nM respectively), while HpARI_CCP1/2 had approximately a 10-fold lower affinity for the cytokine (Kd = 9.8 +/- 6.7 nM). This difference in affinity was largely due to an approximately 20-fold faster off-rate for HpARI_CCP1/2 (K_−_ of 30 x10^-4^ s^-1^ versus 1.5 x10^-4^ s^-1^ for HpARI) (Figure 1A).

**Figure 1:**
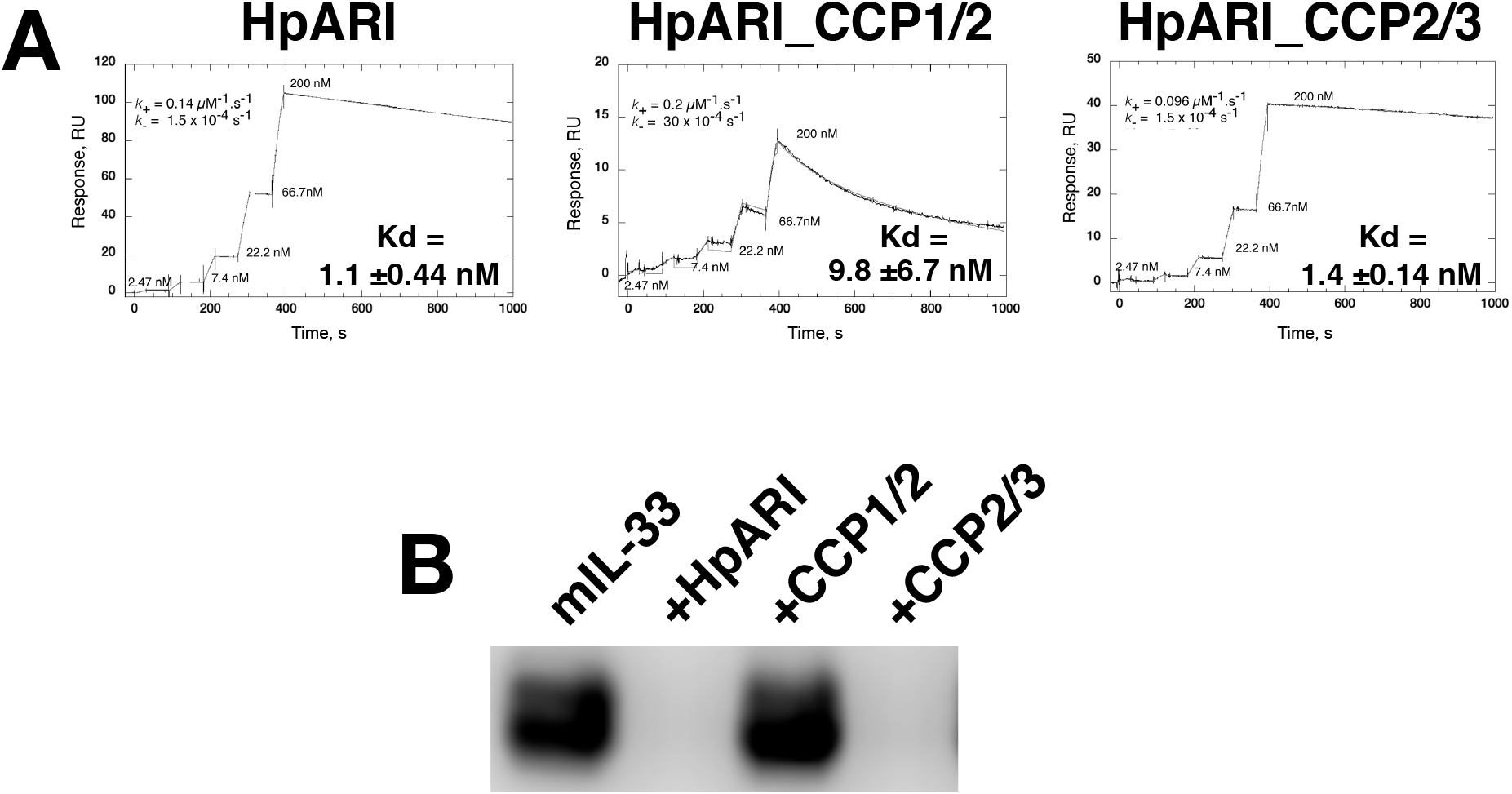
**(A)** Surface plasmon resonance measurements of IL-33 binding to chip-bound HpARI, HpARI_CCP1/2 and HpARI_CCP2/3. **(B)** ST2-Fc was bound to protein G-coated magnetic beads and used to immunoprecipitate murine IL-33 (mIL-33). IL-33 western blot of eluted material shown. Image representative of two independent experiments.

The CCP3 domain also appears important for preventing IL-33-ST2 interactions. While full-length HpARI and HpARI_CCP2/3 were able to prevent IL-33 immunoprecipitation by ST2-Fc, HpARI_CCP1/2 could not (Figure 1B).

### HpARI_CCP1/2 increases responses to IL-33

We previously showed that HpARI_CCP1/2 was capable of suppressing the release of IL-33 in vivo, 15 minutes after *Alternaria alternata* administration (Osbourn et al., 2017). To assess whether HpARI_CCP1/2 could replicate the inhibition of IL-33-dependent responses seen with full-length HpARI, we administered HpARI or HpARI_CCP1/2 together with *Alternaria* allergen and OVA protein and assessed type 2 immune responses after OVA challenge 2 weeks later. While HpARI suppressed allergic reactivity in this model (as shown previously (Osbourn et al., 2017)), HpARI_CCP1/2 had the opposite effect, increasing BAL and lung eosinophil numbers and lung ILC2 numbers (Figure 2A and Supplementary Figure 2).

**Figure 2:**
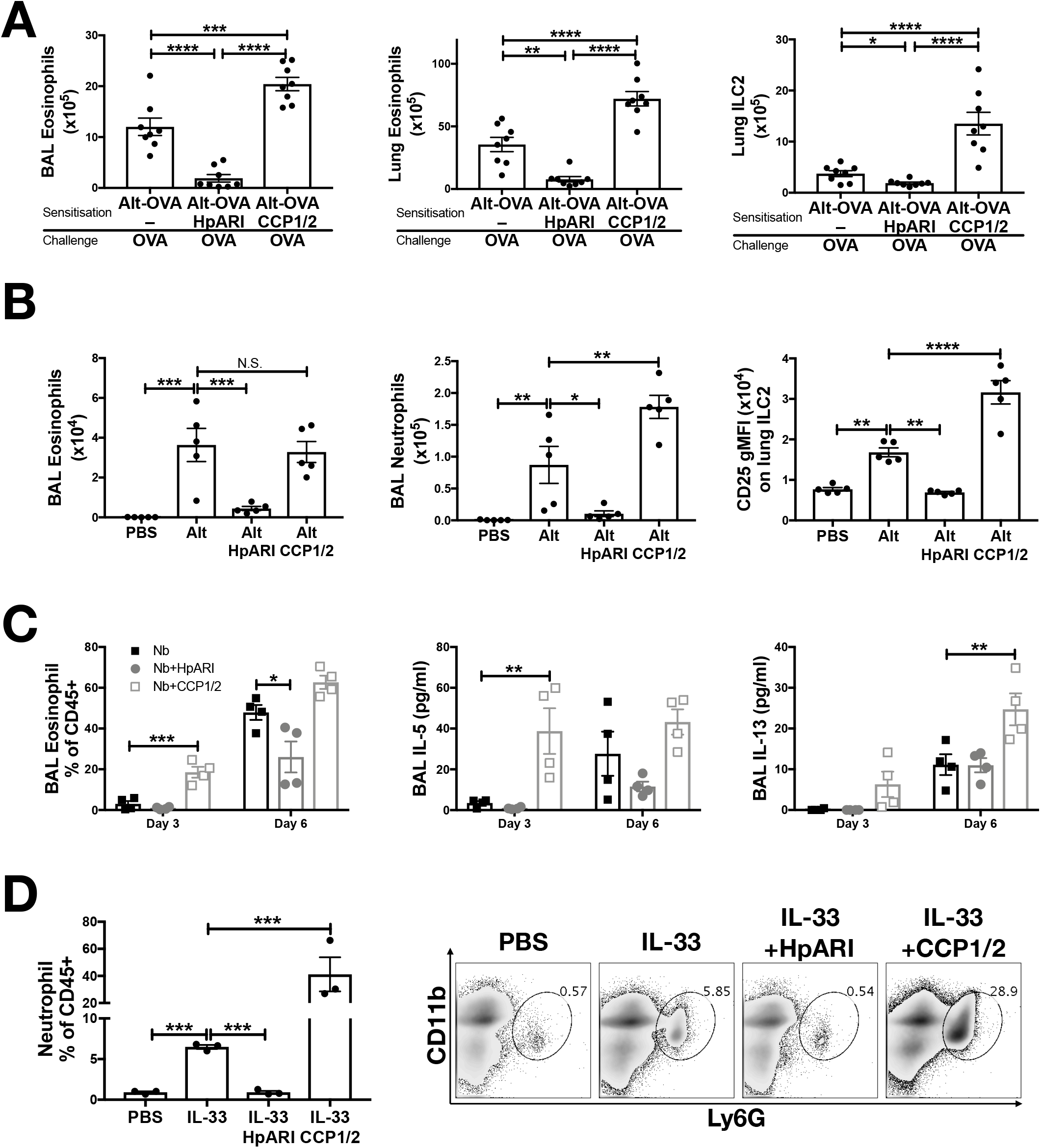
**(A)** HpARI or HpARI_CCP1/2 (CCP1/2) were coadministered with *Alternaria* and OVA by the intranasal route, then the OVA-specific response recalled 2 weeks later. BAL and lung eosinophil (Siglecf+CD11c-CD45+) and ILC2 (ICOS+lineage–CD45+) cell numbers shown. Data pooled from 2 repeat experiments. **(B)** HpARI_CCP1/2 (CCP1/2) was coadministered with *Alternaria* allergen by the intranasal route. After 24 h, BAL eosinophil (Siglecf+CD11c-CD45+) and neutrophil (Ly6G+CD11b+Siglecf-CD11c-CD45+) numbers, and lung ILC2 CD25 geometric mean fluorescent intensity were assessed by flow cytometry. Data representative of 2 repeat experiments. **(C)** HpARI or HpARI_CCP1/2 were intranasally administered on days 0, 1 and 2 after infection with *Nippostrongylus brasiliensis*. BAL eosinophil (Siglecf+CD11c-CD45+) cell numbers, and BAL IL-5 and IL-13 were measured on days 3 and 6 postinfection. Data representative of 3 repeat experiments. **(D)** Recombinant IL-33 was intraperitoneally injected with HpARI or HpARI_CCP1/2, and proportions of Ly6G+CD11b+ neutrophils in the CD45+ peritoneal lavage population assessed 3 h post-injection. Representative FACS plots shown of CD45+ live cells. Data representative of 2 repeat experiments.

Similarly, when the innate *Alternaria-induced* immune response was assessed 24 h after initial administration of the allergen to naïve mice, we found that although HpARI_CCP1/2 did not change the eosinophil response compared to *Alternaria* alone, HpARI_CCP1/2 increased BAL neutrophil numbers. At this timepoint, no ILC2 proliferation has yet occurred (as previously described (Doherty et al., 2012)), so total lung ILC2 cell numbers were similar in all groups (data not shown), however allergen-activated ILC2s showed strong upregulation of CD25 expression (as described previously during activation of ILC2s in this model (Bartemes et al., 2012)), which was further increased by HpARI_CCP1/2 (Figure 2B).

To exclude the possibility that HpARI_CCP1/2 is interfering with the *Alternaria* allergen directly, exacerbating the response to it, we used a second model of IL-33-dependent responses (Filbey et al., 2019; Hung et al., 2013; Wills-Karp et al., 2012), infecting mice with *Nippostrongylus brasiliensis* and administering HpARI or HpARI_CCP1/2 to the lungs during the first 3 days of infection. During *N. brasiliensis* infection, larvae migrate through the lung at days 1-4, and enter as adults into the gut at days 4-10 post-infection (Filbey et al., 2019). Mice were therefore culled at days 3 and 6 post-infection, at the peak of the lung and gut stages, and the type 2 immune response in the lung was assessed. Again, HpARI suppressed type 2 immune responses (as shown previously (Osbourn et al., 2017)), while HpARI_CCP1/2 increased BAL eosinophilia, IL-5 and IL-13 production (Figure 2C).

Finally, we utilised a model of recombinant IL-33 intraperitoneal injection, which induces a mast cell-dependent neutrophilia (Enoksson et al., 2013), in contrast to the ILC2-dependent, largely eosinophilic response seen on IL-33 release in the lung. Again, here we found that while HpARI suppressed IL-33 induced neutrophilia, HpARI_CCP1/2 exacerbated it (Figure 1D).

In conclusion, HpARI_CCP1/2 amplifies IL-33-dependent responses *in vivo*. We hypothesised that this activity was due to stabilisation of the cytokine, increasing its effective half-life. To test this hypothesis, we developed an *in vitro* model of IL-33 release and IL-33 responses.

### HpARI_CCP1/2 maintains IL-33 in its active form

The CMT-64 cell line constitutively produces IL-33, which is released on cellular necrosis (Scott et al., 2018). Confluent CMT-64 cells were washed into PBS+0.1% BSA, and necrosis induced by addition of 0.1% Triton X100, in the presence or absence of HpARI or HpARI_CCP1/2. Over a 24 h timecourse following Triton addition, we assessed IL-33 release by ELISA and western blot. IL-33 ELISA showed that Triton X100 caused rapid IL-33 release, with high concentrations of the cytokine detected in culture supernatants within 15 min of Triton addition in control wells. IL-33 levels then gradually decreased at later timepoints, presumably as the protein was degraded (Figure 3A)(Scott et al., 2018). HpARI addition ablated the IL-33 signal seen in the ELISA, as shown in our previous study (Osbourn et al., 2017): as well as retarding the release of the cytokine, HpARI binding also out-competes the ELISA antibodies, abolishing detection of IL-33. HpARI_CCP1/2 did not abolish detection of IL-33 in the ELISA, but did reduce the IL-33 signal at early timepoints. Moreover, in the presence of HpARI_CCP1/2, IL-33 accumulated over the timecourse and maintained high levels at later timepoints.

**Figure 3:**
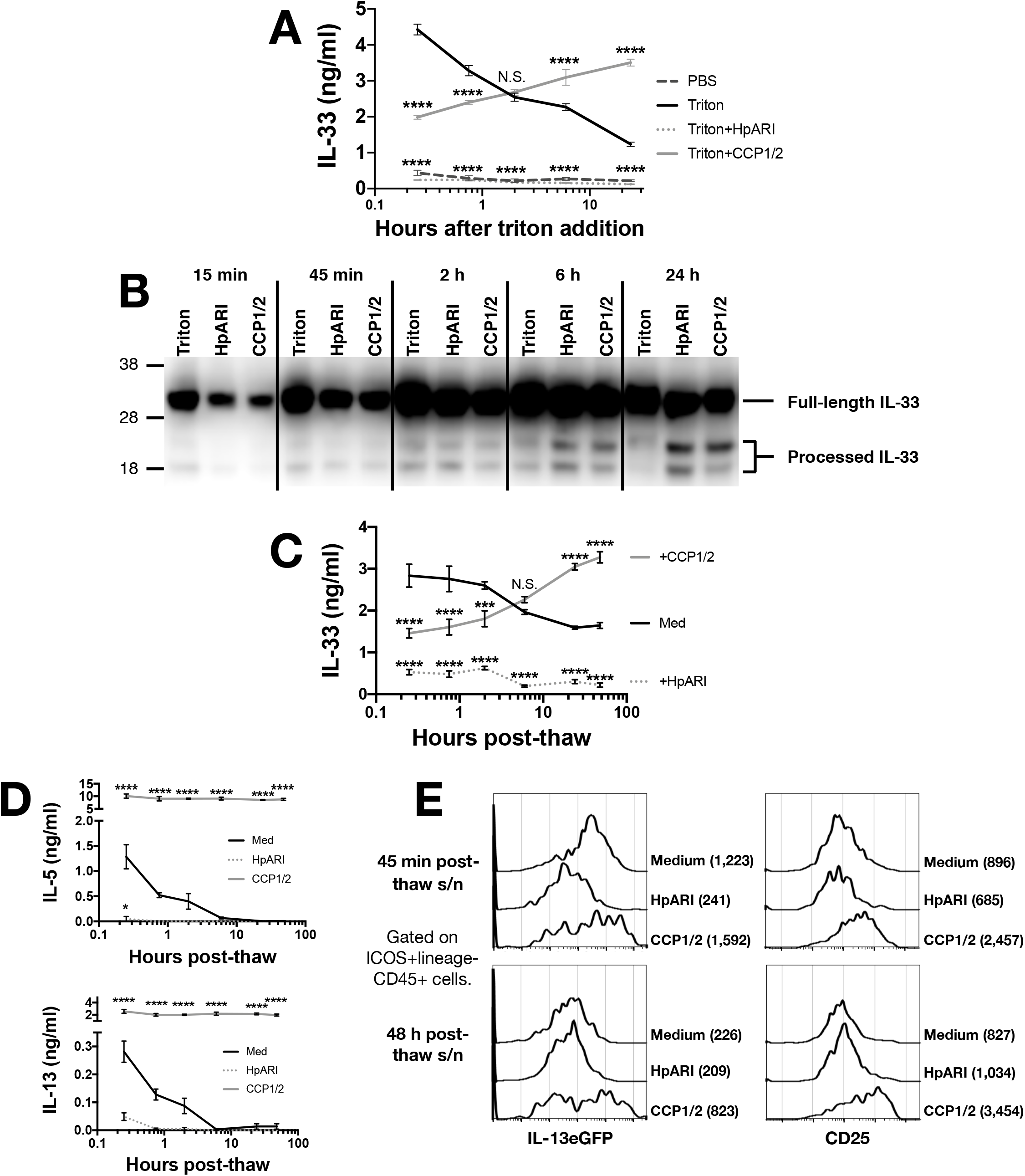
**(A)** CMT-64 cells were cultured to confluency, and treated with 0.1% Triton X100+0.1% BSA alone, or in the presence of HpARI or HpARI_CCP1/2 (CCP1/2). Supernatants were harvested over a timecourse and IL-33 levels assessed by ELISA. Each measurement contains 4 technical replicates and is representative of 3 repeat experiments. **(B)** IL-33 Western blot of pooled samples from (A). Representative of 3 repeat experiments. **(C)** CMT-64 cells were cultured to confluency in RPMI+10% FCS, and freeze-thawed in the presence of complete medium (Med), HpARI or HpARI_CCP1/2. After thaw, cultures of necrotic cells were incubated at 37C, and supernatants taken over a timecourse, and assessed for IL-33 levels by ELISA. Each timepoint shows 4 technical replicates. **(D)** Supernatants from (C) were applied to IL-13eGFP+/- bone marrow cells in the presence of IL-2 and IL-7, and cultured for 5 days. Levels of IL-5 (upper panel) and IL-13 (lower panel) in supernatants were assessed by ELISA. Each timepoint shows 4 technical replicates. **(E)** Bone marrow cells from (D) after 5 days of culture were pooled, stained, and gated on ICOS+lineage-CD45+ ILC2s, and assessed for IL-13eGFP and CD25 expression. Numbers in parentheses indicate geometric mean fluorescent intensity for each condition. All data from C-E is representative of 3 repeat experiments.

In contrast, when IL-33 in the same samples was assessed by western blot, a very strong signal was seen at all timepoints at a size consistent with full-length IL-33 protein (~30 kDa), while a weaker signal was seen at around 18-20 kDa, consistent with processed mature IL-33 (Figure 3B and Supplementary Figure 1). While a strong full-length IL-33 band was seen across all timepoints and treatments, the density of the mature bands were dynamically altered by the presence of each treatment: in control wells, mature IL-33 was present early after Triton-X100 treatment and was degraded at later timepoints, but in the presence of HpARI_CCP1/2, the mature form was present at lower intensities than in control wells at early timepoints, but accumulated over the timecourse and was strongest at 24 h post triton treatment, reflecting ELISA data (Figure 3A). HpARI treatment had a similar effect to HpARI_CCP1/2 when IL-33 was assessed by western blot. The difference in IL-33 signal strength between ELISA and western blot in the presence of HpARI was seen in a previous study (Osbourn et al., 2017), and is thought to be due to interference with antibody binding to the endogenous IL-33-HpARI complex in ELISA, but in a denaturing western blot, proteins from this complex are dissociated and available for antibody detection. Together, this data suggests that binding of IL-33 by HpARI or HpARI_CCP1/2 stabilises the mature cytokine, protecting it from degradation.

To assess the activity of the cytokine released, we induced necrosis of CMT-64 cells via freeze-thaw treatment. This treatment could be carried out in complete culture medium (without toxic additives such as Triton-X100), allowing downstream assessment of cellular responses to the released cytokine. On thaw, necrotic CMT-64 cells were incubated for up to 48 h at 37°C, and IL-33 levels in supernatants assessed by ELISA. Similarly to Triton X100-mediated necrosis, we found high levels of IL-33 released rapidly after freeze-thaw necrosis, which gradually decreased over the 48 h timecourse in control wells, while IL-33 levels increased over the timecourse in the presence of HpARI_CCP1/2 (Figure 3C). These supernatants were applied to bone marrow cells from IL-13^+/eGFP^ reporter mice (Neill et al., 2010) cultured in the presence of IL-2 and IL-7 (to support ILC2 differentiation), and ILC2 responses assessed 5 days later. As shown in Figure 3D, control freeze-thaw CMT-64 supernatants could only induce bone marrow cell IL-5 and IL-13 production at early timepoints post-thaw, implying that after ~6 h post-thaw, all IL-33 present in the culture medium was inactive. This response appeared IL-33-dependent as HpARI entirely inhibited IL-5 and IL-13 release. In contrast, supernatants from cells freeze-thawed in the presence of HpARI_CCP1/2 were able to maintain high levels of IL-5 and IL-13 stimulation (approximately 10-fold higher than the peak production seen in control wells) and this stimulation was maintained even when supernatants had been incubated for 48 h post-thaw. Finally, flow cytometry for IL-13eGFP reporter or CD25 expression was used to confirm that ILC2s were activated by supernatants from medium of freezethaw control wells at early (45 min post-thaw), but not late (48 h post-thaw) timepoints, while wells containing HpARI_CCP1/2 remained highly activated throughout the timecourse (Figure 3E and Supplementary Figure 3).

## 5. Discussion

HpARI blocks IL-33 responses and is secreted by *H. polygyrus*, as part of a suite of immunomodulatory effector molecules which act to prevent immune-mediated ejection of the parasite (Maizels, Smits, & McSorley, 2018). HpARI acts by binding to IL-33 (through the HpARI CCP2 domain) and to genomic DNA in necrotic cells (through the HpARI CCP1 domain), tethering the cytokine within the necrotic cell nucleus and preventing its release (Osbourn et al., 2017). Here, we further characterise these interactions, showing that a synthetic, non-natural construct lacking the CCP3 domain (HpARI_CCP1/2) binds IL-33 with an approximately 10-fold lower affinity than the full-length HpARI protein, and lacks the blocking activity of HpARI against IL-33-ST2 interactions. Furthermore, HpARI_CCP1/2 had the surprising effect of stabilising and amplifying IL-33 responses in vitro and in vivo.

IL-33 is known to mediate parasite expulsion in a type-2 dependent-manner (Hung et al., 2013; Zaiss et al., 2013). The HpARI_CCP1/2 truncated protein maintains the activity of IL-33, potentially amplifying its anti-parasitic effects. It is worthwhile emphasising that this truncated construct is not a protein naturally secreted by the parasite, but rather a synthetic product with an unexpected activity.

As the IL-33 pathway is strongly implicated in human asthma, HpARI, with its unique mechanism of action and strong binding to IL-33, is a potential therapeutic agent. IL-33 is a potently inflammatory cytokine which is kept tightly regulated. Once released, IL-33 undergoes rapid oxidation and degradation, confining its effects to a short time after release (Cohen et al., 2015; Scott et al., 2018). Addition of HpARI or HpARI_CCP1/2 prevented degradation of the cytokine and maintained it in its active form, possibly due to steric hinderance of proteases. As HpARI also blocked the interaction of IL-33 with its receptor, there was no cellular response to IL-33 in the presence of HpARI, while HpARI_CCP1/2, which lacks this IL-33-ST2 blocking activity, was unable to inhibit responses to IL-33. Furthermore, most surprisingly, HpARI_CCP1/2 was able to maintain the effects of IL-33 over a long time-course, potently exacerbating IL-33-dependent responses *in vivo* and *in vitro*.

The effects of HpARI_CCP1/2 may not be confined to extending the half-life of IL-33 by preventing its degradation, but may prevent the much more rapid oxidation of the cytokine. Partial oxidation of IL-33 occurs *in vivo* within 15 min of release (Cohen et al., 2015), therefore the activity of released IL-33 *in vivo* may be less than that of fully active IL-33. Indeed, when a purified wild-type or an oxidation-resistant mutant of human IL-33 were tested *in vitro*, the mutant form of IL-33 was found to be 30-fold more potent than WT IL-33 (Cohen et al., 2015). In this study, we were not able to measure the difference between reduced and oxidised IL-33, therefore we cannot make definitive statements about this activity of HpARI_CCP1/2. However, inhibition of IL-33 inactivation, either through prevention of oxidation or proteolytic degradation, could be a potent method for amplifying IL-33-dependent responses.

Although IL-33 is strongly implicated in inducing eosinophilic inflammation in anti-parasite or allergic type 2 immune responses (Filbey et al., 2019; Liew, Girard, & Turnquist, 2016), the cytokine has also shown protective effects in models of colitis (Lopetuso et al., 2018), graft-versus-host disease (Zhang et al., 2015), autoimmunity (Jiang et al., 2012), obesity (Mahlakoiv et al., 2019), wound healing and tissue restoration (Monticelli et al., 2011; Rak et al., 2016). Therefore, treatments which amplify endogenous IL-33 responses could have clinical potential in a range of treatments.

HpARI_CCP1/2 could also be a useful tool for IL-33 research. Modulating IL-33 responses by using HpARI and HpARI_CCP1/2 in parallel allows assessment of the role of IL-33 in a system in the absence of potentially confounding effects of recombinant cytokine administration or genetic manipulation. In addition, the strategy of IL-33 stabilisation by HpARI_CCP1/2 may be able to be replicated using a monoclonal antibody-based therapy, with low-affinity or non-blocking antibodies potentially able to amplify IL-33 responses. As anti-IL-33 treatments enter clinical trials (Chen et al., 2019), this is an important consideration, as sub-optimal antibodies could result in amplification rather than suppression of IL-33 responses.

This study sheds further light on the mechanism of binding of HpARI to IL-33, the function of the domains of HpARI, and the effects of IL-33 degradation and inactivation. Further structural characterisation of HpARI–IL-33 binding will be useful in characterising this interaction and could allow guided design of more effective IL-33-blocking or IL-33-amplifying therapeutic agents.

## Acknowledgements

Flow cytometry data was generated with support from the QMRI Flow Cytometry and cell sorting facility, University of Edinburgh.

## Author Contributions

CC, FV, MW and HM designed and planned experiments. CC, FV, SC, JR, WG, AO, MW and HM undertook experiments. MW provided guidance on the design of the SPR experiments and carried these out. HM supervised all work and wrote the paper.

---

**Supplementary Figure 1:**
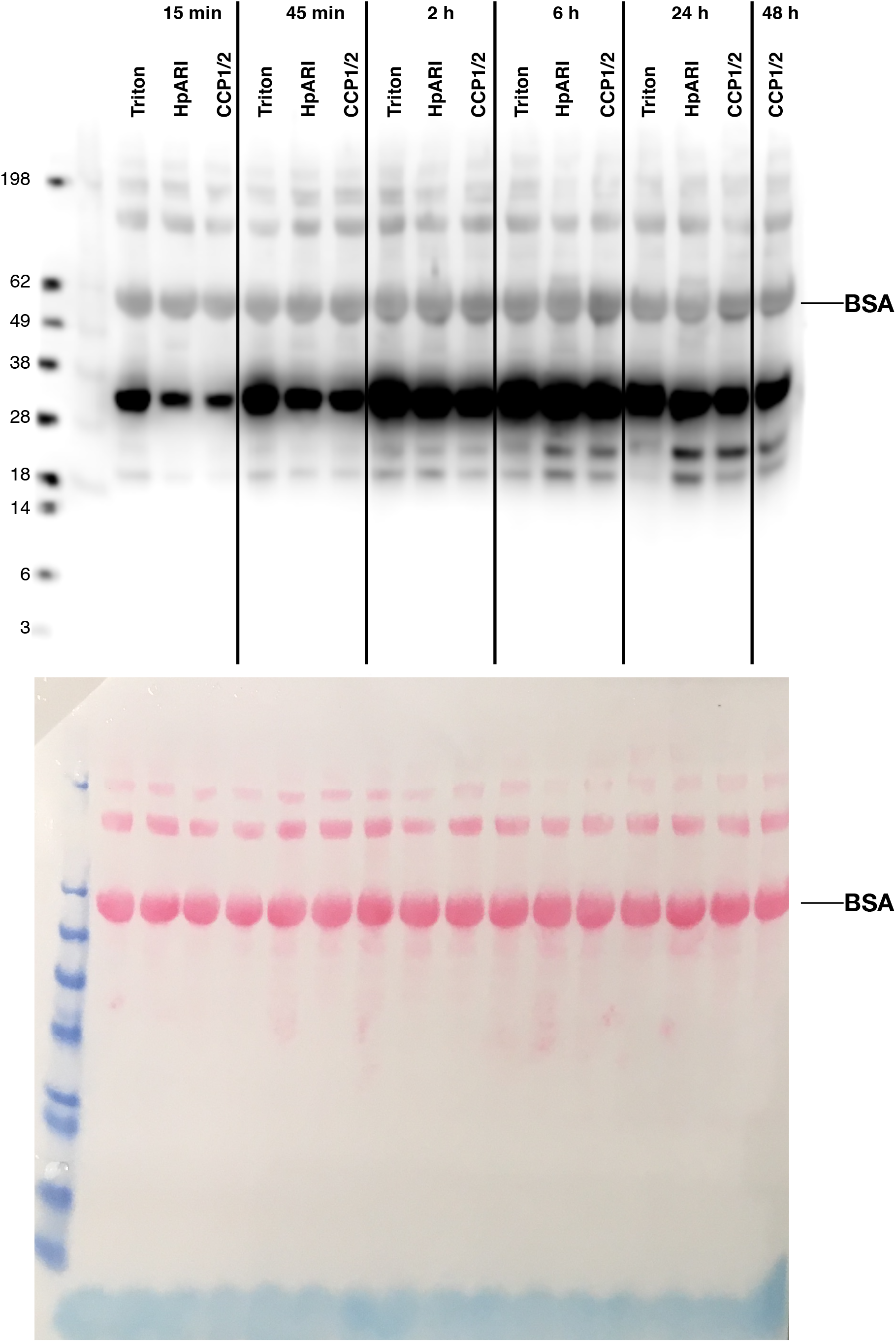
**(A)** Uncropped western blot shown in Figure 3B. **(B)** Ponceau stain of blot shown in (A)

**Supplementary Figure 2:**
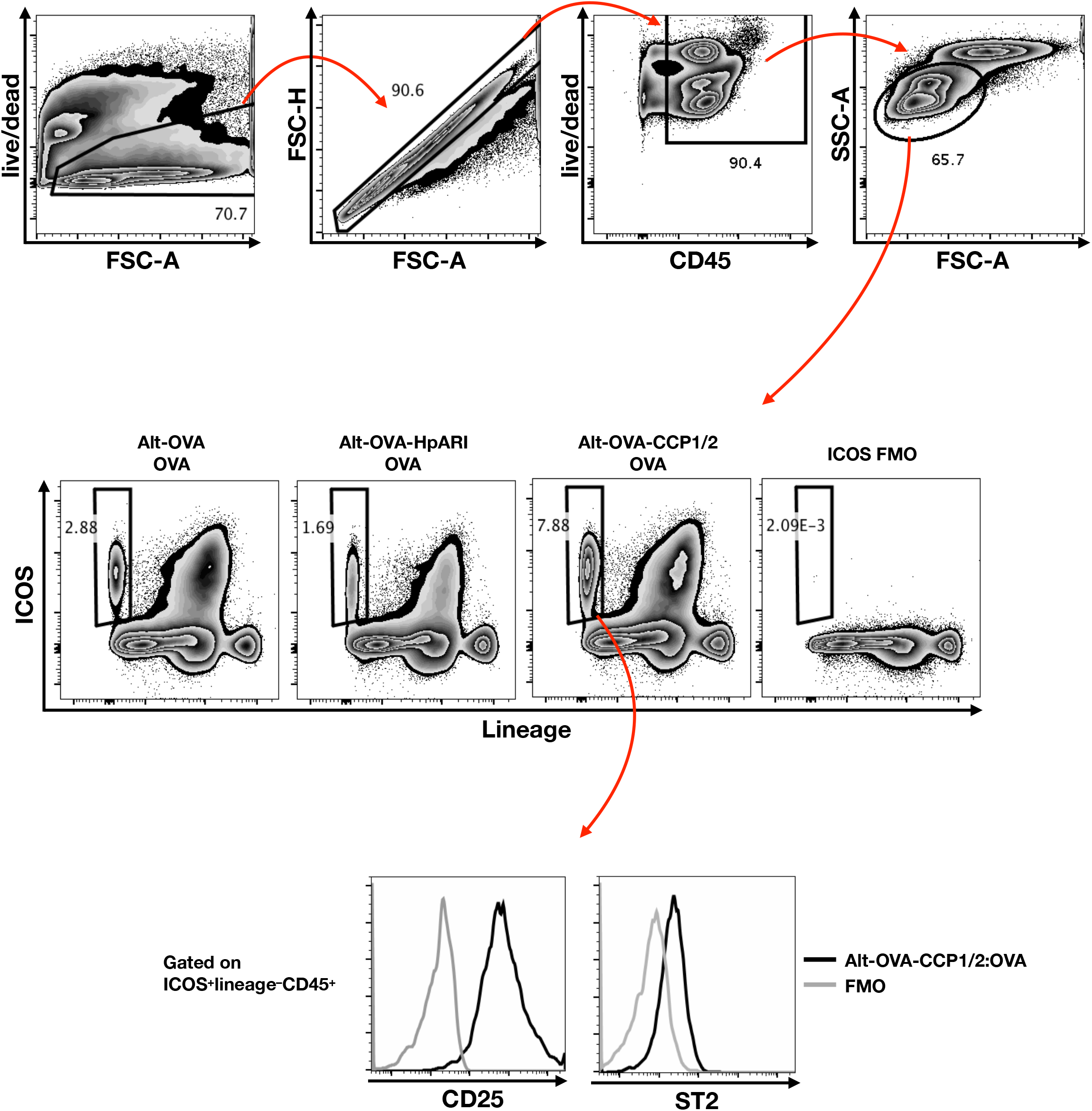
Gating strategy and FMO controls for lung innate lymphoid cell staining, from samples described in Figure 2A.

**Supplementary Figure 3:**
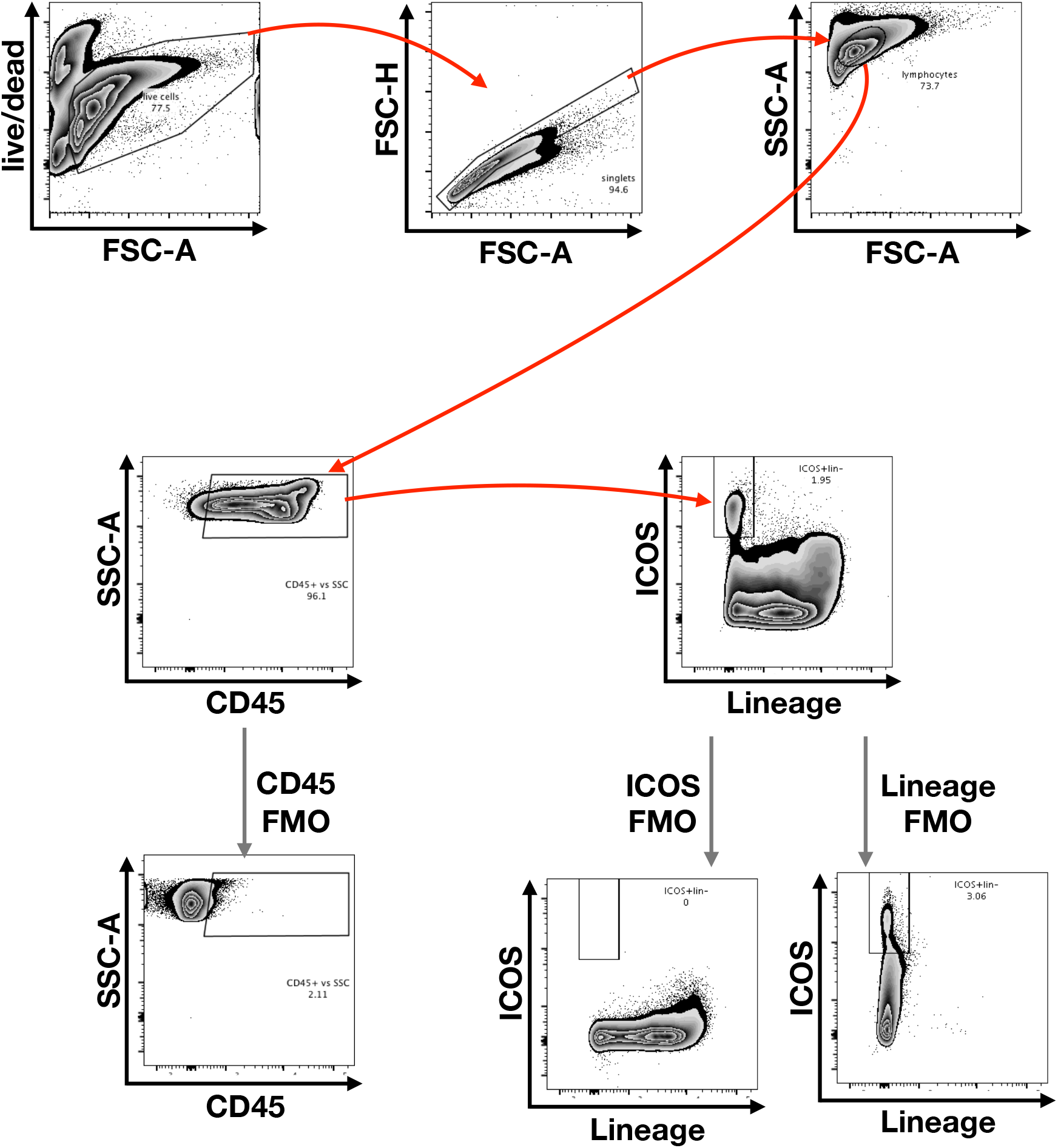
Gating strategy and FMO controls for ICOS+lineage–CD45+ ILC2s shown in Figure 3E

